# Forging Sustainable Research: Integrating Sustainability into Neuroscience Education

**DOI:** 10.64898/2026.10.05.756692

**Authors:** William V. Smith, Aimee Bebbington, Hannah Bowker, Tessa Christbaum, Stefan R. Pulver

## Abstract

The health and wellbeing of living organisms and the environment are irrevocably interconnected. Animals constantly shape the environments in which they live, and environmental factors, in turn, shape the minds and behaviours of animals. The emerging discipline of environmental neuroscience studies this reciprocal interaction and offers a unique opportunity to integrate sustainability into undergraduate training. Here, we present and evaluate ‘*Forging Sustainable Research’*, an interdisciplinary undergraduate course that integrates teaching of neuroscience, biology, psychology, environmental science, and sustainable development. Students explore how research in these fields can be interlinked and re-oriented towards addressing emerging environmental challenges. In parallel, students learn how to measure and mitigate the environmental impact of research activities. Student feedback suggests that the course is effective at raising awareness and understanding of sustainability in research, while also enhancing student perception of their own ability to promote positive change. As climate and ecosystem change impact all sectors, sustainability awareness in academic and professional spaces is increasingly sought after. Students in the course therefore receive training in transferrable skills relevant for diverse career paths. In the long term, this course may serve as model for direct integration of sustainability training into undergraduate curricula in the natural sciences.

## Introduction

The environment constantly and unavoidably shapes the neurobiology and behaviour of living organisms. Anthropogenic climate change is rapidly altering ecosystems (Intergovernmental Panel On Climate Change (IPCC), 2023): global rising temperatures and the bioaccumulation of waste products are straining ecosystems and consequently altering behaviours of animals that live within them (Michaiel & Bernard, 2022). The growing field of environmental neuroscience – the study of how the environment shapes nervous systems and behaviour, and vice versa – highlights how little we understand about how rapidly-changing ecosystems are altering the behaviour and health of living organisms (Brierley & Kingsford, 2009; Chevance et al., 2023; Kühn & Gallinat, 2023; Rocque et al., 2021). From altering the neural networks of crustaceans (Brierley & Kingsford, 2009; Marder & Rue, 2021) to differential effects on neurological disease (Berntsson et al., 2025; Jilanee et al., 2025; Louis et al., 2023; Martínez Lozada & Leon-Rojas, 2025; Reis et al., 2024; Sisodiya et al., 2024), climate change is fundamentally shaping the neural basis of behaviour.

Climate education is not simply an exercise in knowledge transfer, but involves shaping future competency. As the effects of climate change quickly impact different sectors of society, knowledge of sustainability in academic and professional spaces is becoming a sought-after competency (Bilderback, 2024; Redman & Wiek, 2021; Smith et al., 2025). Education thus represents a key anchor for forging a sustainable future (UNESCO, 2020). Despite the urgent need for sustainability competency, there exists a significant, global gap in mainstream climate education (QAA, 2021; UNESCO, 2020). While there exist strong efforts in discipline-integrated sustainability education, particularly in the form of short online courses (Angstmann et al., 2019; Dunne et al., 2022; Freese et al., 2024; Sikand et al., 2021; Wolfe & Lindeborg, 2018), to our knowledge there are no reported environmental neuroscience modules which integrate quantitative carbon accounting and action-orientated sustainability training. Many pedagogical approaches to sustainability education have difficulty bridging the knowledge-action gap (Bowen et al., 2025; Tang, 2022). Consequently, there is a need for teaching approaches that empower the citizens of tomorrow to understand, reflect on, and act justly in the spirit of good environmental policy both within and beyond academia. Given the tangible ways climate change influences behaviour and nervous systems, integrating climate and neuroscience education offers a valuable opportunity to train the next generation of citizens and researchers in key sustainability-related competencies (Wolfe & Lindeborg, 2018).

Here, we present and evaluate an interdisciplinary undergraduate course centred on environmental neuroscience and sustainability-related sciences. Across a semester, final-year neuroscience undergraduates explored topics in environmental neuroscience and neuroethology; examined the sustainability of common neuroscience and other research techniques; studied the evolving landscape of sustainability within and beyond academia, and gained hands-on experience in key sustainability competencies including life cycle emissions assessments and communication of environmental neuroscience research. Students reported significantly higher awareness of key sustainability concepts and increased confidence after completing the course. Our work shows how neuroscience and sustainability can be directly integrated to bridge knowledge-action gaps, improving graduating students’ sustainability literacy and developing increasingly in-demand skills in carbon accounting.

## Materials and Methods

### Curriculum & Teaching Methodology

The one-semester course (“Forging Sustainable Research”) was offered to final-year neuroscience, biology, and psychology undergraduate students at the University of St Andrews. Students (N=10) were introduced to a variety of topics related to environmental neuroscience and sustainability through hour-long sessions twice a week for 12 weeks (Table 1). Each week was typically composed of a lecture and discussion session focused on developing connections between neuroscience theory and research techniques and their environmental consequences. Within these discussion sessions, a group of students would present a pre-selected paper to commence the discussion, guided by the demonstrator. For instance, the course examined the diversity of neuroscience research tools (e.g., optogenetics (Häusser, 2014)), the ways in which those tools are being used at the frontiers of research, and the environmental costs of different stages of their life cycle (i.e., production, usage, and disposal-related emissions). Within each topic, a follow-on student-led discussion session predominately centred on a pre-determined publication (see below). This was used to reinforce the content of earlier lectures and provide a space for critical thinking and ethical discussion about the environmental consequences of research.

**Table 1:**
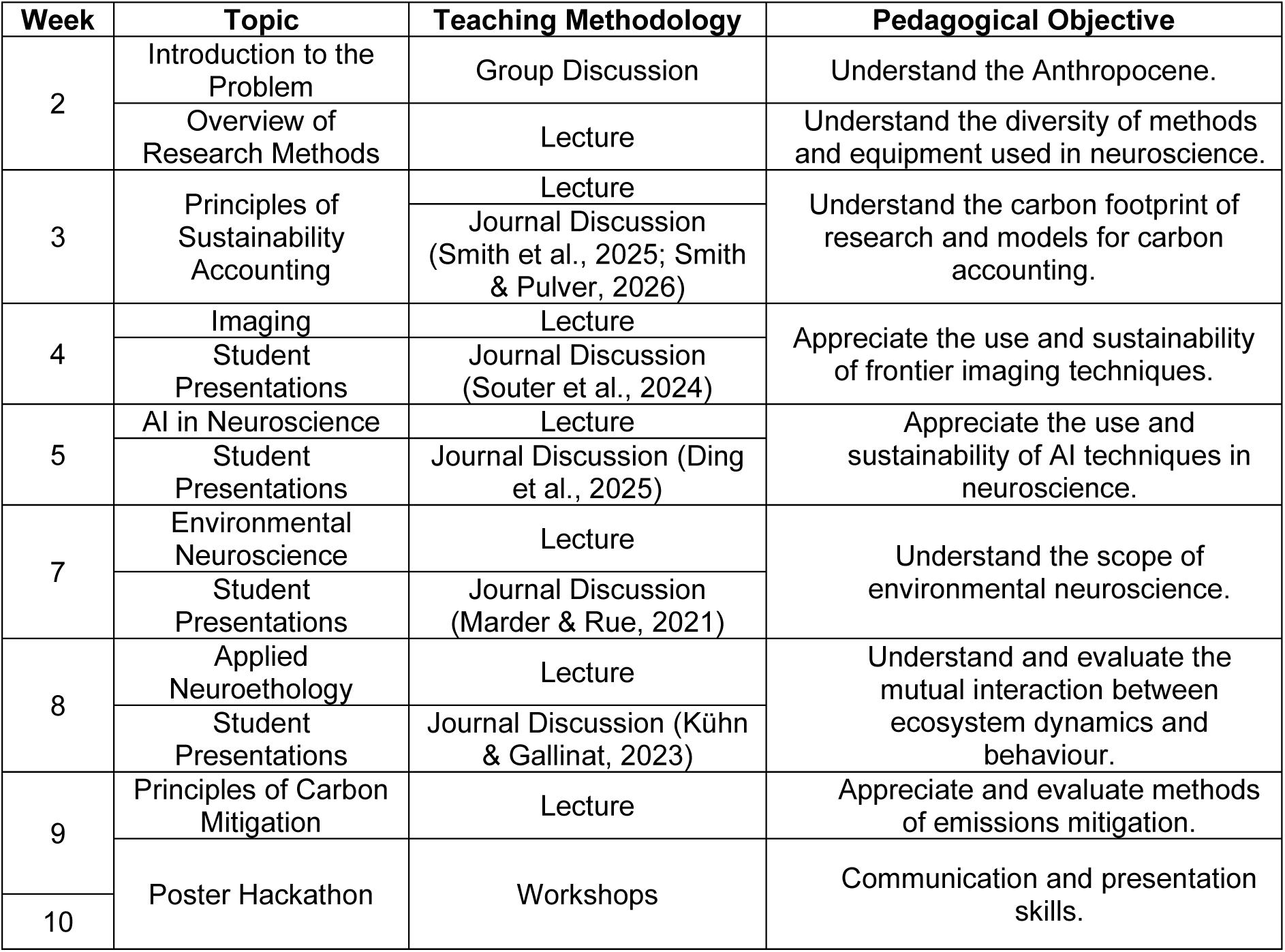
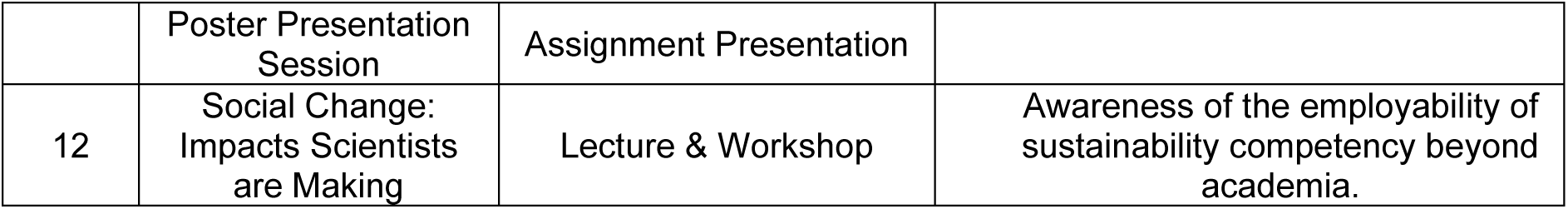
Course topics, teaching methodology, and pedagogical objectives.

Amongst the rising abundance of literature on environmental neuroscience and climate change (Rae et al., 2022; Sisodiya, 2026; Todorova et al., 2025), the topics in the module were selected to balance and synergise the expertise of the instructors with current and rising sustainability demands in academia and beyond. The topic profiles were grounded in the expertise of the teaching staff, who have expertise in neuroethology, neurobiology, computational neuroscience, ecology, and carbon accounting. Further, the selected topics introduced students to the sustainability implications of common neuroscience research methods(e.g.., imaging), as well as presenting a forward-focused and beyond-academic selection of ideas, such as artificial intelligence and social change. Additional literature beyond Table 1 is referenced within the lecture notes and often arose during free-flowing class discussions. An important goal of the module was to provide a broad perspective of ways in which individuals are shaping sustainability across society (Dupont et al., 2024; Racimo et al., 2022; Urai & Kelly, 2023). Within a ‘Social Change’ session in the final week of the course, we invited guest speakers from beyond academia to discuss their experience with sustainability. Previously, we hosted a representative from a commercial bank who discussed how major UK banks have begun offering financial incentives on mortgages to improve the energy performance of homes, as well as tree-planting scheme for new-build gardens (Fatouh et al., 2025). Providing an external perspective on sustainability enabled students to ground their understanding of climate action beyond academia and appreciate the transferable benefits of sustainability knowledge, training, and skills.

### Assessments

The course was exclusively assessed through two coursework components: a procurement report (50%), and an environmental neuroscience poster (50%). The topics of both coursework assessments were chosen by the undergraduate students with the guidance of demonstrators. Exemplar reports of varying grade classification were offered to the students. General task instructions and a marking rubric were also supplied tostudents. The course followed the University of St Andrews generative AI policy which forbids the use of generative AI in the generation or marking of coursework material except under reasonable adjustments approved by the Disability Support Team (Academic Policy Officer, 2026).

#### Procurement Report

The first assessment required students to create a procurement report for an item of research. The purpose of the procurement report is to present an estimate of the carbon dioxide equivalent emissions associated with procurement (i.e., scope 3 emissions) of a research-related item (for example, a specific chemical reagent, or a pipette tip) (Intergovernmental Panel On Climate Change (Ipcc), 2023b). To calculate scope 3 emissions, students were required to select an item of importance to research and contact a supplier of that item; trace the supply chain from that supplier to St Andrews; and report an estimate of the resulting scope 3 emissions based on distance calculations, item weight, and reported UK Government emissions factors (Smith et al., 2025). The students were given the freedom to select any item relevant to neuroscience research, with part of the procurement assessment evaluating the student’s justification for why they selected their item. By design, students were often confronted with an absence of complete information about their chosen item’s supply route which necessitated making and justifying educated guesses of scope 3 emissions. Students were required to reach out to companies for supply-related information as part of the assessment. Evidence of writing to an external partner was required, but final marks did not depend on the student receiving a response. Across 8 procurement reports, 7 for-profit manufacturers or suppliers were engaged, alongside 3 non-profit research institutions (some students contacted multiple sources).

The procurement report assessment consolidated key topic points within the course, including understanding of research methodologies, emission calculations, problems of information transparency, and the effect of research methodology on emissions. Critically, the assessment empowered the students to develop carbon literacy and accounting skills while also engaging directly with industry partners. The freedom of item choice and the necessity to justify item choice and methodology all encourage personal tailoring of the coursework to achieve maximal engagement, utility, and thus pedagogical outcome. Within their report, students were expected to comment on the item’s utility within common experimental designs with the very best reports discussinghow usage rates would affect scope 3 emissions. Stronger submissions were differentiated based on students’ commentary on the product’s larger life-cycle, and their originality in proposing ways to mitigate the environmental impact of its procurement beyond changing provider. An example of a summary figure presented in a procurement report is found in the results section (Figure 1).

**Figure 1:**
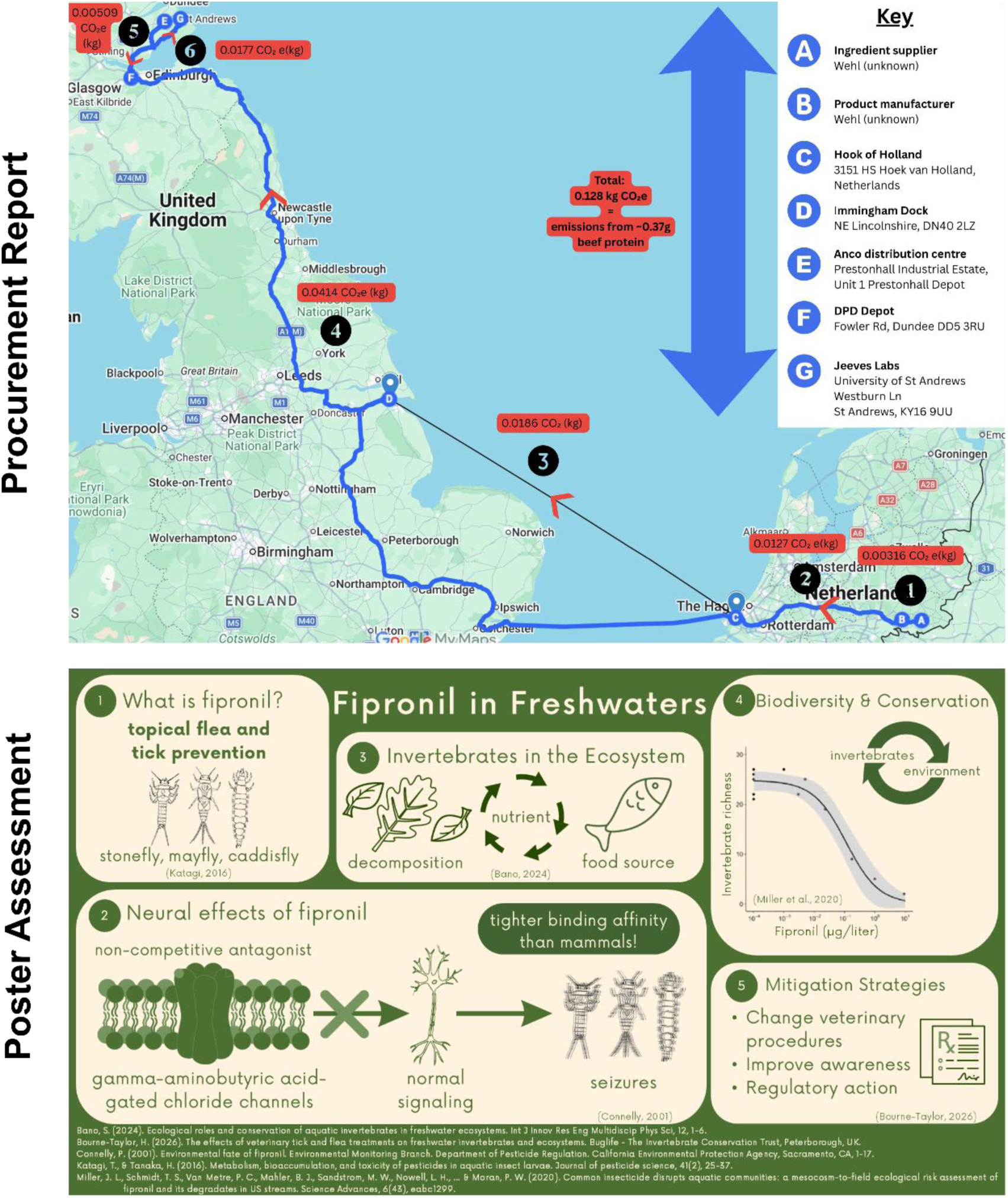
Examples of Student work. Example of a procurement report summary figure and a student’s environmental neuroscience poster. The procurement report exemplar demonstrates the foundation of the procurement report – contextualising scope 3 emissions estimates based on transportation of real-world scientific items. In this report, the student estimated emissions arising from procurement of food for canine research at the University of St Andrews by considering either ocean freight or heavy-goods vehicle transportation routes. The procurement assessment expects students to choose an item of importance to scientific research, justify the necessity for a procurement-related emissions estimate based on product usage rates, then calculate relative scope 3 emissions based on real-world information about supplier location, travel routes, and distance-to-emissions factors reported by government or private agencies. The exemplar environmental neuroscience poster demonstrates one student’s approach to this assessment, namely identification of an environmental stressor and its known effects on living organisms. The student evaluated the effects of the broad-spectrum phenylpyrazole insecticide Fipronil on freshwater invertebrate movement health, and explored broader implications for biodiversity stability and methods for mitigation of its environmental impacts.

#### Environmental Neuroscience Poster

The second assessment required students to create and present a poster on a topic in environmental neuroscience. The field of environmental neuroscience is broad and rapidly evolving (Kühn & Gallinat, 2023). Students were encouraged to choose a focused topic that explores how environmental factors shape neurobiological processes and behaviour, and vice versa. A wide variety of appropriate topics were presented ranging from molecular environmental neuroscience (e.g., the effects of insecticide pollution in freshwater systems on invertebrate and vertebrate behaviour) to ethology (e.g., the impact of biodiversity change on the hunting behaviour of wolves) to human psychology (e.g., the relationship between the natural environment and human health). To aid students in creating their posters, dedicated hackathon sessions were placed into the teaching schedule to give students the opportunity to ask questions, integrate feedback, and discuss the role of environment in shaping neurobiological processes and behaviour.

### Student Survey

All research followed ethical guidelines, with the student survey receiving approval by the internal ethics committee of the School of Psychology & Neuroscience at the University of St Andrews. Students were invited to take part in surveys at the beginning (week 2) and end (week 12) of the course. Surveys were hosted on Qualtrics with anonymous links distributed by QR codes within the class and through reminder emails after the first and final weeks of teaching. Students self-generated anonymous IDs to compare before and after surveys. The survey was explicitly advertised as being entirely optional andparticipation anonymous. 55% of students enrolled in the course completed both instances of the survey.

The survey predominately examined changes in students’ knowledge and perceptions as a result of participating in the course. Each instalment of the survey consisted of 7 questions to assess changes in students’ knowledge and perspective:

1. How aware and knowledgeable are you on the following sustainability topics: environmental sustainability research funding, radiative forcing, carbon load of academic research, methods for carbon accounting, AI in neuroscience, green research programs (e.g., LEAF, MyGreenLabs), environmental neuroscience, sustainable agriculture? *Unaware (0); Aware (1.25); Knowledgeable (2.5); Experienced (3.75); Expert (5)*.
2. What is your perspective on being able to overcome the following challenges in sustainability: research sustainability, green transition, climate change acceptance, climate change? *Apathetic (0); Neutral (3); Optimistic (5)*.
3. How much power and influence do you think the following people or institutions have on affecting sustainability: students, universities, funding bodies, companies, governments, international bodies, financial sector? *None at all (0); A little (1.25); A moderate amount (2.5); a lot (3.75); a great deal (5)*.
4. What words come to mind when you think of research sustainability?
5. How likely are you to include attempts to improve sustainability in your future professional life? *Extremely Unlikely (0); Somewhat Unlikely (1.25); Neither Likely nor Unlikely (2.5); Somewhat Likely (3.75); Extremely Likely (5)*.
6. How much influence do you think you as a student have to improve sustainability? *None at all (0); A little (1.25); A moderate amount (2.5); a lot (3.75); a great deal (5)*.
7. What is the main limitation you feel hampers your ability to engage in sustainability: knowledge, time, motivation, outcome likelihood, resources (e.g., tools), professional support (e.g., teachers, managers)? (1 - most, 6 - least).

### Statistical Analysis of Survey Data

Anonymous data was analysed and visualised using a Jupyter Python notebook. We performed statistical comparisons of students’ awareness and perspectives before and after teaching. Changes in reported awareness or perspective are declared as relative difference scores (d). Normality was assessed using a Shapiro-Wilk test. In instances of normality, a paired t-test was performed. In instances of violation of normality, a Wilcoxon signed-rank test was performed. Reported p-values were corrected across comparisons using the Benjamini– Hochberg false discovery rate procedure.

## Results

### Course operation and activities

We ran *Forging Sustainable Research* over the course of 2 years with a formal student survey conducted in year 2. In both years, the course ran largely as planned with minor changes made to scheduling and content – primarily to increase discussion of relevant current events. The course had one lead instructor (SP) organizing most lectures and discussions but also incorporated guest lectures and workshops from other faculty and postgraduates.

### Assessment Engagement

Engagement with both the procurement report and environmental neuroscience poster assessment was generally high. A diverse array of research item procurement emissions estimates was documented, and environmental neuroscience topics explored (Table 2). Notable exemplars of a procurement report summary figure and an environmental neuroscience poster are shown in Figure 1.

**Table 2:** Range of Procurement Assessment Focuses.

| Topics |  |
| --- | --- |
| <u>Environmental Neuroscience</u> | <u>Procurement Focus</u> |
| Mercury Exposure to Cognitive Deficits | Sharpie Pens |
| Behavioural Neuroscience of Yellowstone Wolf Hunts | Fish Feed |
| Fipronil in Freshwater | Guinea Fowl Eggs |
| Temperature Effects on Turtle Sex Ratio | Dog Food |
| Virtual Nature Improving Mental Health | Dry Ice |
| Effects of CO<sub>2</sub> on *R. maidis* Foraging and Feeding Behaviour | Foam Earplugs |
| Effect of Anthropogenic Noise on Marine Mammal Stress and Reproduction | Signagel® Electrode Gel |
| BMMA Accumulation Research and Finning Prohibition | Leica Microsystems Cryostat Research Instrument |

### Teaching Outcomes

Students’ anonymous reporting of topic awareness revealed a noticeable improvement in understanding across the major topic themes in the course (Figure 2). Notably, students (N=6) reported becoming significantly more aware of and knowledgeable about the carbon footprint of research (d=2.7, p=.0495), environmental neuroscience (d=3.0, p=.02), methods of carbon accounting (d=2.8, p=.03), research funding aimed at improving sustainability (d=1.7, p=.03), and the rise of sustainability auditing initiatives like LEAF and MyGreenLabs (Schell & Bruns, 2024) (d=1.2, p=.03).

**Figure 2:**
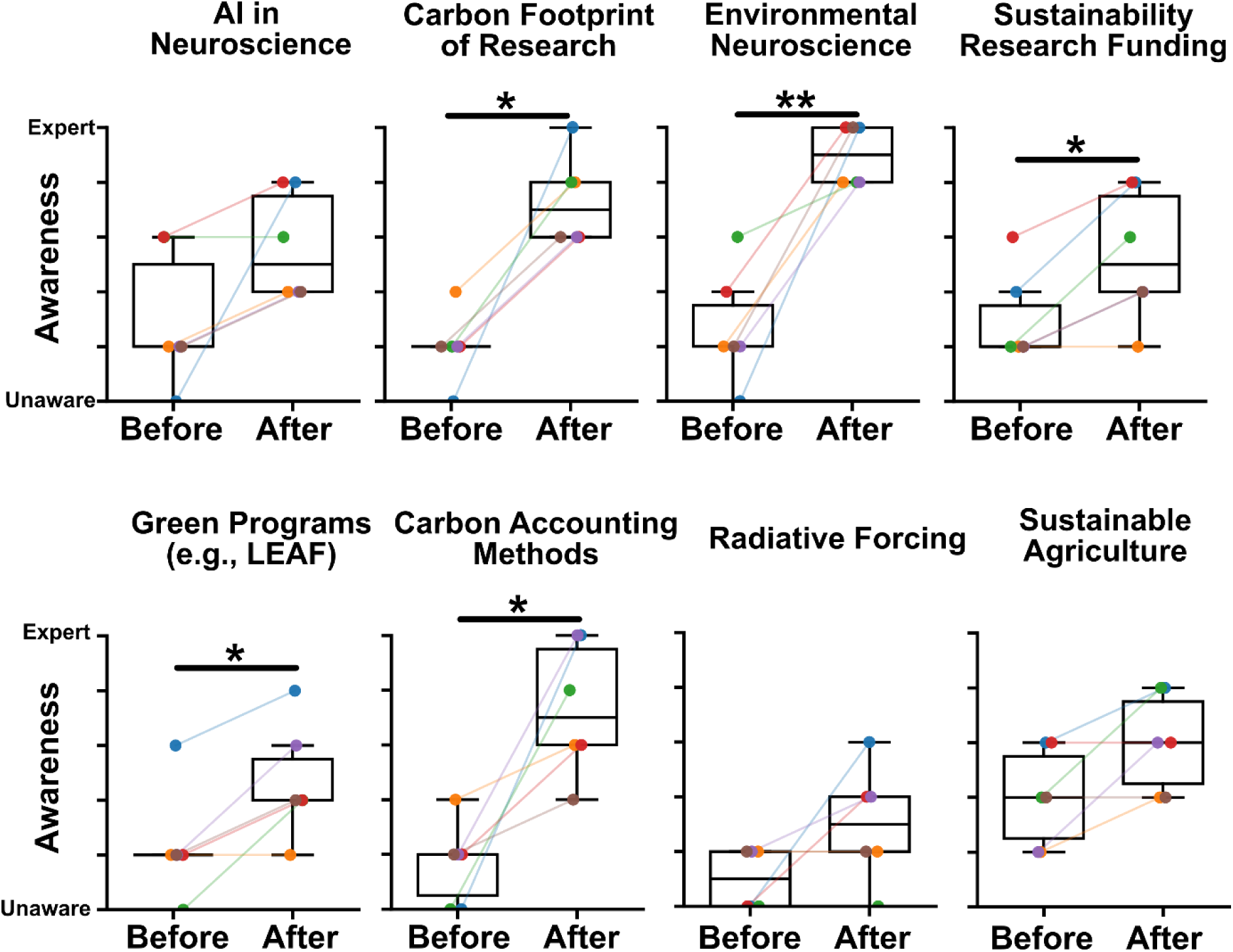
Students showed increased awareness and understanding of sustainability in research during the course. Reported changes in awareness (0 = unaware, 5 = expert) from surveyed students (N=6) of key teaching topics.

### Students’ Changed Perspectives

Students reported positive shifts in recognition of their own role for positively influencing sustainability. While teaching did not significantly alter students’ perspectives on the being able to overcome climate-related challenges (Figure 3A), the course significantly improved students’ perspective on their level of power and influence in positively shaping sustainability (d=1, p=.0495). Interestingly, students generally thought most large-scale bodies (i.e., individual companies, universities, governments, and funding bodies) have a high level of power to shape sustainability (Figure 3B). By the end of the course students associated a greater variety of terms to sustainability, with a shift away from conceptions of sustainability through ‘waste’ and ‘reduction’ towards a greater emphasis on being ‘conscious’ and technical terminology like ‘CO_2_e’ (Figure 3C). Further, students reported a consistently high likelihood of including efforts to improve sustainability in their future career (Figure 3D). Interestingly, the students’ self-reported perspectives on their own personal ability to influence sustainability did not change by the end of the course despite, as show in Figure 3B, the cohort generally perceived students to have higher influence on improving sustainability than they had initially reported at the start of the module (Figure 3E). Finally, while there was individual variability in how each student conceptualised the greatest limiting factor to their own capacity to improve sustainability (Figure 3Fi), generally knowledge, resources and tools remained their top-ranked perceived limiter (Figure 3Fii).

**Figure 3:**
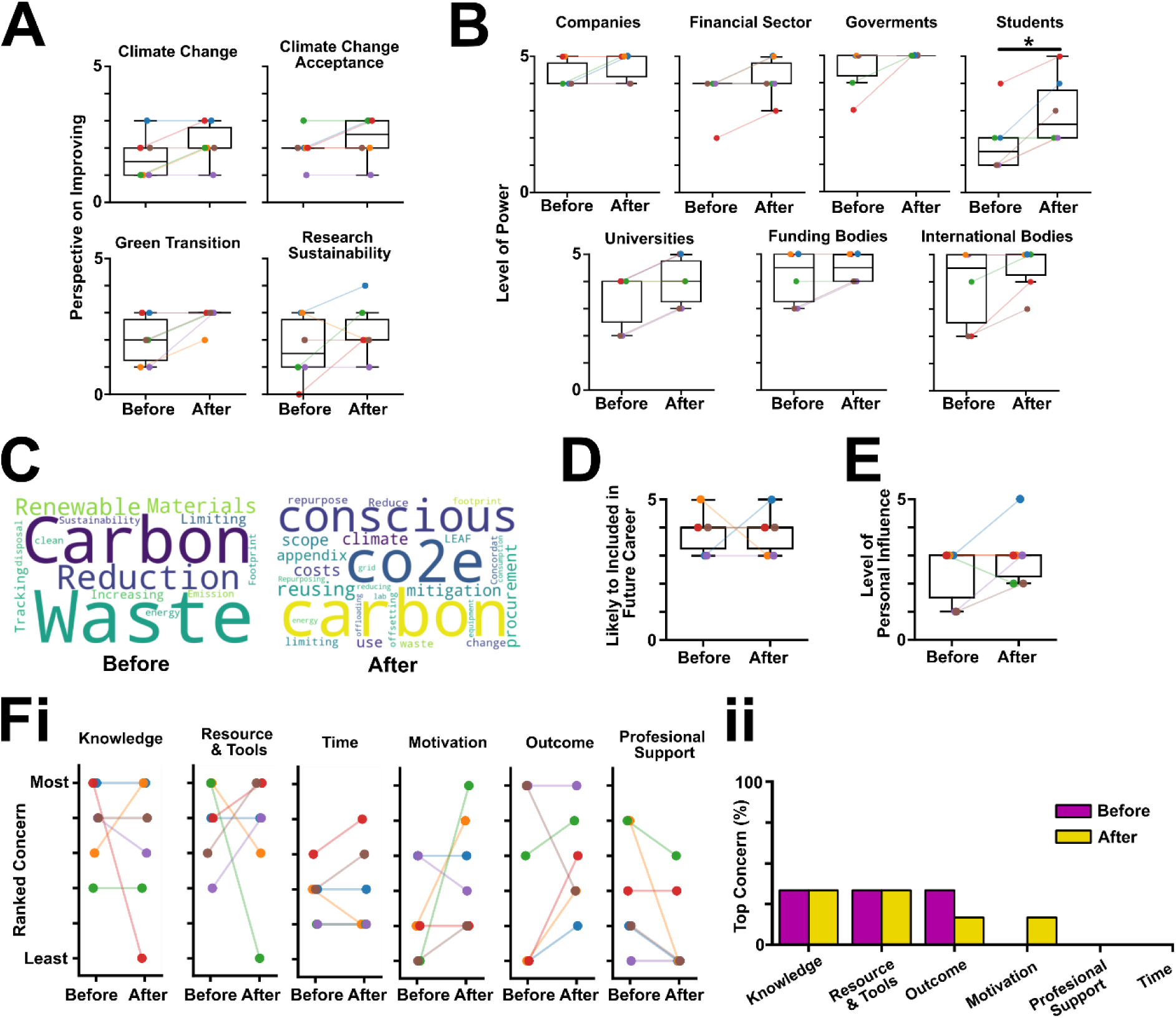
Changed Perceptions and Reflections on Sustainability. (**A**) Reported changes in students’ perspective of being able to overcome societal challenges (0 = apathetic, 5 = optimistic). (**B**) Reported changes in students’ perceived power and influence over the sustainability of people or institutions (0 = none at all, 5 = a great deal). (**C**) Words associated with research sustainability before and after completion of the course. (**D**) Reported changes in likelihood of including attempts at improving sustainability in students’ future professional life (0 = extremely unlikely, 5 = extremely likely). (**E**) Reported changes in perceptions of the level of influence students have to improve sustainability (0 = none at all, 5 = a great deal). (**F**) Reported changes in the perceived main limitation that hampers students’ engagement with sustainability (**i**) across dimensions and (**ii**) percentage of students ranking each dimension as their main limitation (top concern) before and after completion of the course.

## Discussion and Conclusion

At first glance, neuroscience and environmental sustainability can appear to be very different topics. However, closer examination reveals many points of intersection between the two fields. Environmental factors profoundly shape the neurology and behaviour of both invertebrates (Bilderback, 2024; Goncharov et al., 2023; Lawrence & Soame, 2004; Martins Medeiros & Souza, 2023; Pearce-Higgins & Morris, 2022; Shi & Li, 2024) and vertebrates (Anttonen et al., 2023; Michaiel & Bernard, 2022; Seebacher & Little, 2026), including humans (Martínez Lozada & Leon-Rojas, 2025; Reis et al., 2024; Sisodiya et al., 2024). Recent research has begun to reveal the neural mechanisms underlying these changes (Marder & Rue, 2021; Michaiel & Bernard, 2022; O’Donnell, 2018; Robertson & Money, 2012; Schapiro & Marder, 2024; Sisodiya et al., 2024). Conversely, the behaviours of animals, and humans in particular, are driving rapid changes in ecosystems in every part of the world (Hautier et al., 2015; He & Silliman, 2019; Prakash & Verma, 2022). The psychological processes underlying human decision making in this pivotal and turbulent era are increasingly under scrutiny. Environmental neuroscience is emerging as a focal point that brings together interdisciplinary research in these areas that can catalyse action beyond academia.

When students perceive large gaps between knowledge and capability for action, apathy can set in (Fallis, 1991). Here, we show how students can be synergistically introduced to emerging and established neuroscience techniques, concepts in environmental neuroscience, and topics in sustainable development. Further, we demonstrate how coursework focused on forging sustainability competencies and direct engagement with industry can positively engender a understanding of sustainability as well as an increased sense of influence and power to enact positive change. In line with our previous work on sustainability (Smith et al., 2025; Smith & Pulver, 2026), we believe that a pedological approach that centres and empowers students and early-career researchers to become changemakers by allowing them creative freedom to engage with sustainability in their work is an essential part of forging a sustainable future. Our procurement assessment demonstrates how students can simultaneously learn the theory and challenges of greenhouse gas emissions estimation while contributing behaviour-shaping product-specific emissions data. Rapidly changing ecosystems are affecting human and animal behaviour in ways we have yet to fully comprehend (Kühn & Gallinat, 2023). By integrating sustainability into diverse tertiary education curricula, teachers and students can positively shape the future of not just academia, but all spheres of influence.

Education remains relevant by meeting the needs, desires, and interests of its students. Student self-reporting data demonstrated a significant change in awareness and understanding across some topics in the course (e.g., environmental research, but not AI in neuroscience). Further, while students broadly perceived ‘student’ influence being greater as a result of the teaching, they did not attribute this change to themselves as individuals. Indeed, students appeared not to be become any more optimistic about our capacity to remedy climate change by the end of the course. While there is a well-documented low correlation between knowledge gain within education and impactful action (Alam et al., 2025), the *way* students are taught may be integral to the promotion of future sustainable behaviour. As shown here, expanding teaching beyond information delivery towards open discussions and skills-based assessments, alongside a persistent focus on the progress individual students are making towards improving sustainability is one potential avenue to address the intention-behaviour gap (Fielding & Hornsey, 2026; Kukowski et al., 2026). Evaluation of the effects of instructional design on the intention-behaviour gap may inform adaptation of sustainability pedagogy. Future courses may evaluate how students’ perceptions and careers develop post-education, and how companies change their operations in response to interactions like those conducted for our procurement assessment. . Accurate information delivery alone may be insufficient to achieve lasting positive behaviour change. Actively engaging diverse students in sustainable action while communicating the positive impacts on sustainability of other individuals’ actions may engender greater positive outcomes (Holz & Tuitjer, 2026; Pearce et al., 2015; Suldovsky, 2017).

One way to improve engagement with sustainability efforts is to integrate sustainability education into core elements of the student experience. A drawback of our investigation is that this course is an elective in the last semester of a 4-year degree. Sustainability-interested students may be more likely to opt into taking our module, resulting in a cohort that is initially more aware of the challenges faced by climate change, and perhaps, more pessimistic about the power of individuals to effect positive change. Future work could investigate methods to tailor sustainability education to a broader cohort in 1^st^ and 2^nd^ year compulsory modules, and critically evaluate changed student knowledge and perspectives in consequence. Interested educators could include sustainability as a theme across their core lectures. For instance, teachers could assign optional reading about the climate implications of AI (Chouksey et al., 2026; Cowls et al., 2023; Pimenow et al., 2024) or discuss the lack of publicly available information around industrial manufacturing processes and procurement, as well as how researchers are individually improving the sustainability of chemistry research (Kirchhoff, 2005; Sheldon, 2017, 2018). There are sustainability dimensions to nearly every aspect and discipline of research. Strong pedagogical approaches are centred on consistency and repetition but are also focused on actively engaging the student (Luo et al., 2026; Santos et al., 2019). Educators may consider which topics and pedagogies are most likely to encourage positive and active engagement by their students. For instance, open discussion and coursework that yields novel results may encourage student engagement more effectively than traditional lecture-style communication of information. Furthermore, short courses such as lab-based modules could be adapted to incorporate key sustainability considerations and skills, such as industry sustainability standards (Bosi et al., 2022) or lab sustainability auditing using LEAF and My Green Lab (Schell & Bruns, 2024), thereby helping to build students’ awareness of sustainability and transferable skills. However, there will always remain the trade-off between academics’ time, student interest, and the ability to make a genuine impact. Consequently, academics would be wise to consider approaching interested students and tailoring teaching materials to their interests and needs.

Sustainability and climate change are unavoidable concerns not only for neuroscientists and educators, but increasingly for all professions. ‘*Forging Sustainable Research*’ provides a blueprint for harnessing the enthusiasm of societies’ future changemakers to address and improve sustainability, irrespective of their role or profession.

## Acknowledgements

We thank the students and staff participating in PN 4112 *Forging Sustainable Research*. This project was made possible by an Industrial CASE PhD studentship (UKRI Biotechnology and Biological Sciences Research Council (BBSRC) grant number BB/T00875X/1) to WVS, by a Scotland’s Future Series award to WVS, and a World Leading PhD Studentship (University of St Andrews) to A.B., M. Bischoff and J. Kursawe.

## Data Availability

Exemplar course material, marking rubrics, and lecture notes can be found in an open-access Zenodo repository at: https://doi.org/10.5281/zenodo.21923325.

## Address correspondence to

William V. Smith, School of Psychology & Neuroscience, University of St Andrews.

Dr. Stefan R. Pulver, School of Psychology & Neuroscience, University of St Andrews.

